# Practical unidentifiability of receptor density in target mediated drug disposition models can lead to over-interpretation of drug concentration data

**DOI:** 10.1101/123240

**Authors:** Andrew M Stein

## Abstract

For monoclonal antibodies, mathematical models of target mediated drug disposition (TMDD) are often fit to data in order to estimate key physiological parameters of the system. These parameter estimates can then be used to support drug development by assisting with the assessment of whether the target is druggable and what the first in human dose should be. The TMDD model is almost always over-parameterized given the available data, resulting in the practical unidentifiability of some of the model parameters, including the target receptor density. In particular, when only PK data is available, the receptor density is almost always practically unidentifiable. However, because practical identifiability is not regularly assessed, incorrect interpretation of model fits to the data can be made. This issue is illustrated using two case studies from the literature.

## Introduction

When characterizing the pharmacokinetics of monoclonal antibodies (mAbs), mathematical models of target mediated drug disposition (see the Full Model and its Michaelis-Menten approximation in Figure 1) are often fit to preclinical and clinical data in order to estimate key physiological parameters of the system. These parameters (listed in Table 1) can then be used to build a deeper understanding of the underlying physiology, which can guide predictions about the druggability of the target and the minimally effective dose for first in human trials [1, 2, 3].

**Figure 1:**
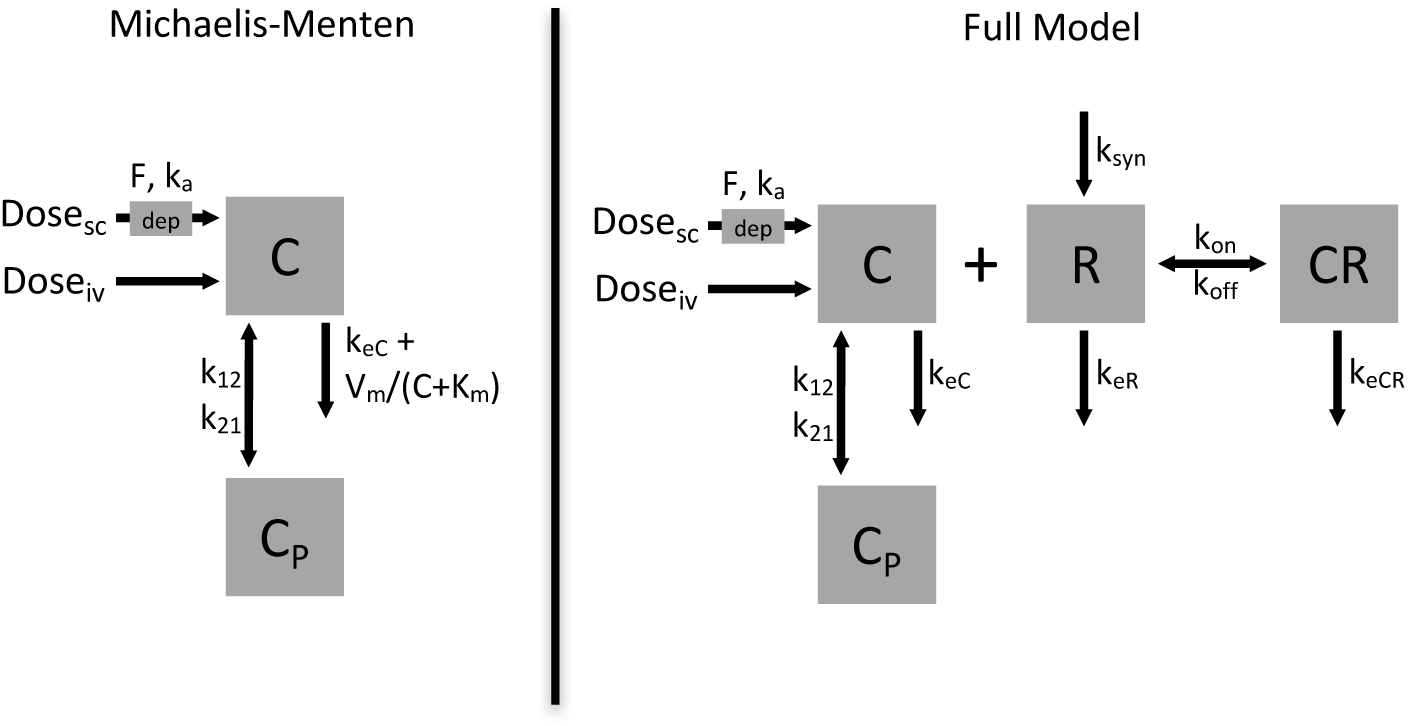
The Michaelis-Menten model and the full target mediated drug disposition model (TMDD) for characterizing monoclonal antibody pharmacokinetics. Parameter descriptions are provided in Table 1.

**Table 1:** Model parameter descriptions.

| Parameter | Type | Description | Units |
| --- | --- | --- | --- |
| $In_{sc}(t)$ | Amount | Subcutaneous dosing function | nmol |
| $In_{iv}(t)$ | Amount | Intravenous dosing function | nmol |
| $C$ | Conc. | free drug concentration | nM |
| $R$ | Conc. | free receptor concentration | nM |
| $(CR)$ | Conc. | complex concentration | nM |
| $C_{tot}$ | Conc. | total drug conc. = $C + (CR)$ | nM |
| $R_{tot}$ | Conc. | total receptor conc. = $C + (CR)$ | nM |
| $k_a$ | Dose | subcutaneous absorption rate | 1/d |
| $F$ | Dose | subcutaneous bioavailability | - |
| $V_c$ | PK | central volume | L |
| $k_{eC}$ | PK | drug elimination rate = $CL/V_c$ | 1/d |
| $k_{12}$ | PK | central → peripheral distribution | 1/d |
| $k_{21}$ | PK | peripheral → central distribution | 1/d |
| $CL$ | PK | clearance = $k_{eC} \cdot V_c$ | L/d |
| $Q$ | PK | inter-compartmental clearance = $k_{12}V_c$ | L/d |
| $V_p$ | PK | peripheral volume = $k_{12}V_c/k_{21}$ | L |
| $k_{syn}$ | Receptor | receptor synthesis rate | nM/d |
| $V_m$ | Receptor/PK | maximum saturable elimination rate $\approx k_{syn}V_c$ | nmol/d |
| $k_{eR}$ | Receptor | receptor elimination rate | 1/d |
| $k_{eCR}$ | Receptor | complex elimination rate | 1/d |
| $R_0$ | Receptor | initial receptor concentration = $k_{syn}/k_{eR}$ | nM |
| $R_{tot,ss}$ | Receptor | steady state total receptor conc. after large dose = $k_{syn}/k_{eCR}$ | nM |
| $k_{off}$ | Binding | disassociation rate | 1/d |
| $k_{on}$ | Binding | association rate | 1/(nM d) |
| $K_{ss}$ | Binding | quasi-steady-state (QSS) const = $(k_{off} + k_{eCR})/k_{on}$ | nM |
| $K_m$ | Binding/PK | Concentration of half saturable elimination for MM | nM |

A challenge in fitting these models to data is that although these models have been shown to be structurally identifiable [4], they are often practically unidentifiable. A model is defined to be structurally identifiable (also referred to as a priori or theoretically identifiable) if there is a one-to-one mapping between the model parameters and the measured variables (e.g. drug concentration). For a structurally identifiable model, it is possible for the investigator to collect sufficient data to estimate all model parameters as long as the assay is of sufficient sensitivity and a range of dosing regimens can be explored. A model parameter is defined to be practically identifiable (also referred to as deterministic, a posteriori, or numerically identifiable) with respect to a particular experimental design; a model parameter is practically identifiable if the confidence intervals on that parameter are finite on a log-scale [5]; i.e. the lower confidence limit of a parameter must be greater than zero and the upper confidence limit must be less than infinity. A non-zero lower limit is required because there can be a big difference between a binding affinity of 1 pM, 1 nM and 1 *μ*M.

An example of a practically identifiable system is when one fits an Emax model *E*(*C*) = *E*_max_*C*/(*EC*_50_ + *C*) to a limited dataset. If only low concentration data is available such that *E*(*C*) looks linear, then E_max_ will be practically unidentifiable with with no upper bound. On the other hand, if only large concentration data is available such that the measured effect ranges from 70%-100% of *E*_max_, there may not be sufficient data to identify a lower bound for the EC_50_, making EC_50_ practically identifiable.

For target mediated drug disposition models for mAbs, the data that is collected is often rich, with intravenous and subcutaneous dosing over a 100x dose range; however, the data is almost never rich enough to identify all model parameters. In particular, PK sampling just after intravenous dosing is usually not frequent enough to estimate the rate of binding (*k*_on_). The assay is also often not sensitive enough to estimate the receptor density (*R*_0_) or drug-receptor complex internalization rate (*k*_eCR_). The mathematical analysis from Peletier and Gabrielsson [6] highlights the conditions under which each parameter is identifiable for a one-compartment model though to our knowledge, an analysis of the two-compartment system has not been published.

One example of practical unidentifiability that is well understood is that even when the binding rate (*k*_on_) and unbinding rate (*k*_off_) are difficult to estimate, a lumped parameter such as *K*_*d*_ = *k*_off_/*k*_on_ or *K*_ss_ = (*k*_off_ + *k*_eCR_)/*k*_on_ may still be identifiable. Thus the pharmacometrics community makes frequent use of the quasi-equilibrium or quasi-steady-state approximations [7, 8].

Another example of practical unidentifiability that is less widely understood is that for membrane-bound targets, the data is often not rich enough to allow for identification of the receptor density at baseline and steady state (*R*_0_, *R*_tot,ss_) and the bound and unbound receptor elimination rates (*k*_eCR_, *k*_eR_). Instead, only lumped parameters are identifiable such as *V*_*m*_ ≈ *k*_syn_ · *V*_c_ = *k*_eCR_ · *R*_tot,ss_ · *V*_c_ and *K*_m_ ≈ *K*_ss_ · (*R*_tot,ss_/*R*_0_). While this issue has been explored in the literature [8, 9], the pharmacometrics community does not regularly employ this insight. In particular, two recent manuscripts make use of TMDD models to estimate the baseline receptor density (*R*_0_) to support physiological understanding of the system for romosozumab [10] or to make PK predictions in humans based on cynomolgus monkey data [11]. However, as will be shown here, *R*_0_ was practically unidentifiable in both instances and only an upper bound for *R*_0_ could be identified.

## Methods

### Models

The following models, derived previously in [8, 12] were explored in this analysis:

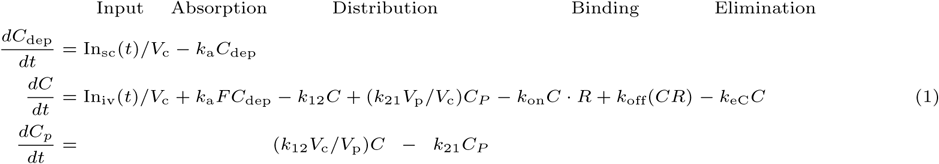

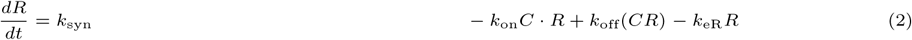

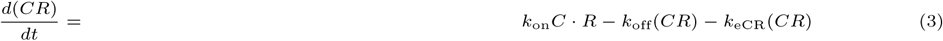

Following a single IV bolus dose, the initial conditions for this model are *C*_dep_(0) = *C_p_*(0) = *CR*(0) = 0, *C*(0) = Dose/V_c_ and *R*(0) = *k*_syn_/*k*_eR_.

### Quasi-steady-state model (QSS)

For drugs with membrane-bound targets, binding and complex internalization generally happen quickly and it is assumed that RC is in a quasi-steady-state such that

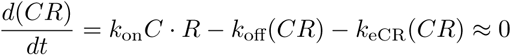

Rearranging the above equations for *C*·*R*/(*CR*) gives the equation below, where now the differential equation above for *RC* has been replaced with this algebraic expression.

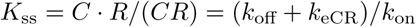

The differential equations can then be written in terms of total drug and receptor levels giving:

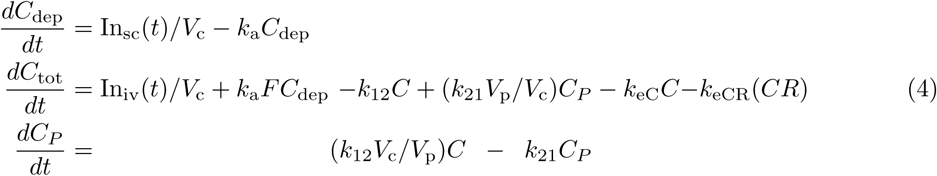

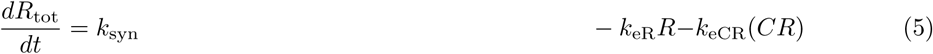

The initial conditions following a single IV bolus dose are *C*_dep_(0) = *C*_*P*_ (0) = 0, *C*_tot_(0) = Dose/*V*_c_ and *R*_tot_(0) = *k*_syn_/*k*_eR_. The concentrations *C, R*, and (*CR*) on the right hand side are given by the equations below, which were derived by substituting *C* = *C*_tot_ − (*CR*) and *R* = *R*_tot_ − (*CR*) into the equilibrium equations above and solving for (*CR*).

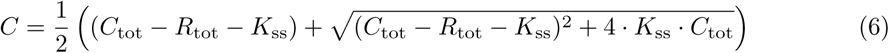

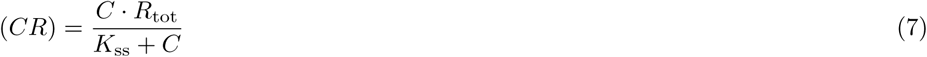

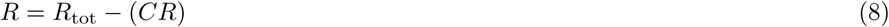

### Constant Turnover (CT)

The constant turnover approximation assumes that *k*_eCR_ = *k*_eR_ and thus that *R*_tot_ is constant. A method was developed for replacing the differential equations and algebraic expressions for the equilibrium models above with a single differential equation [13, 9], though in the simulations performed here, the constant turnover assumption is applied simply by setting *k*_eCR_ = *k*_eR_ in the QSS model.

### Michaelis-Menten (MM)

The Michaelis-Menten approximation is given by:

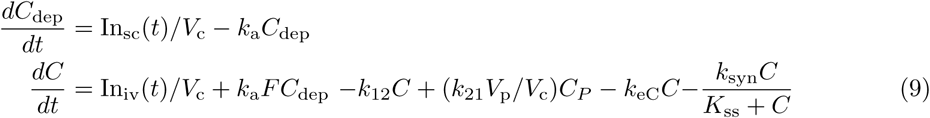

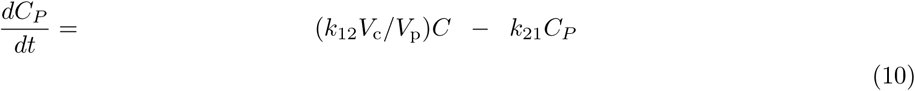

The initial conditions following a single IV bolus dose are *C*_dep_(0) = *C*_P_ (0) = 0 and *C*(0) = Dose/*V*_c_.

### Estimating the baseline and large-dose receptor number

Consider Equation 5. When no drug is in the system, C = (CR) = 0 and

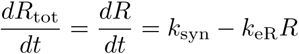

At steady state when *dR*_tot_/*dt* = *dR*/*dt* = 0 gives *R*_0_ = *k*_syn_/*k*_eR_ in the absence of drug. For large doses, almost all drug is bound, giving *R*_tot_ ≈ (*CR*) and *dR*_tot_/*dt* ≈ *d*(*CR*)/*dt*. Thus:

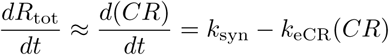

At steady state, *d*(*CR*)/*dt* = 0, giving *R*_tot,ss_ ≈ (*CR*)_ss_ = *k*_syn_/*k*_eCR_ following large doses. This was previously demonstrated by Peletier and Gabrielsson [6].

### Sensitivity analysis

To explore a larger range of models and parameter values and to build intuition for how the parameters impact the PK profile, a sensitivity analysis was undertaken for the Michaelis-Menten, QSS-CT, QSS, and full model. In this analysis, one parameter was varied while all other parameters were held fixed. The parameters explored were: {dose, *V*_m_, *CL,V*_c_, *V*_p_, *Q, K*_m_, *R*_tot,ss_,*R*_0_, *k*_off_}. To perform this sensitivity analysis in such a way that one lumped parameter could vary while the others were held fixed, the other rate constants are calculated in terms of the lumped parameters, as shown below.

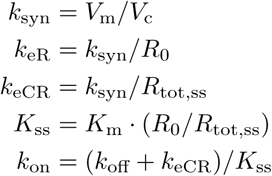

In this analysis, the equation *K*_m_ = *K*_ss_ · (*R*_tot,ss_/*R*_0_) was used. This relationship was chosen because it was found that varying *K*_ss_ · (*R*_tot,ss_/*R*_0_) for the QSS and full model gave comparable behavior to varying Km in the Michaelis-Menten model, as shown in the Results section. This product of an equilibrium constant times the ratio of receptor density at steady state to baseline also appeared in [14], where the AFIR metric (average free target to initial target concentration) was defined for mAbs binding soluble targets. There, *K*_d_ was used instead of *K*_ss_ because the elimination rate of the drug-target complex is much slower for mAbs with soluble targets. Note that when the constant turnover approximation is employed, *R*_tot,ss_ = *R*_0_ and *K*_m_ = *K*_ss_. The parameters used in the sensitivity analysis are provided in the caption of Figure 2 and were based on the fits to the mavrilimumab sensitivity analysis. Because it will be shown that *R*_0_ is unidentifiable from the mavrilimumab data, a large range of *R*_0_ and *R*_tot,ss_ is explored, from 0.01 nM to 100 nM.

**Figure 2:**
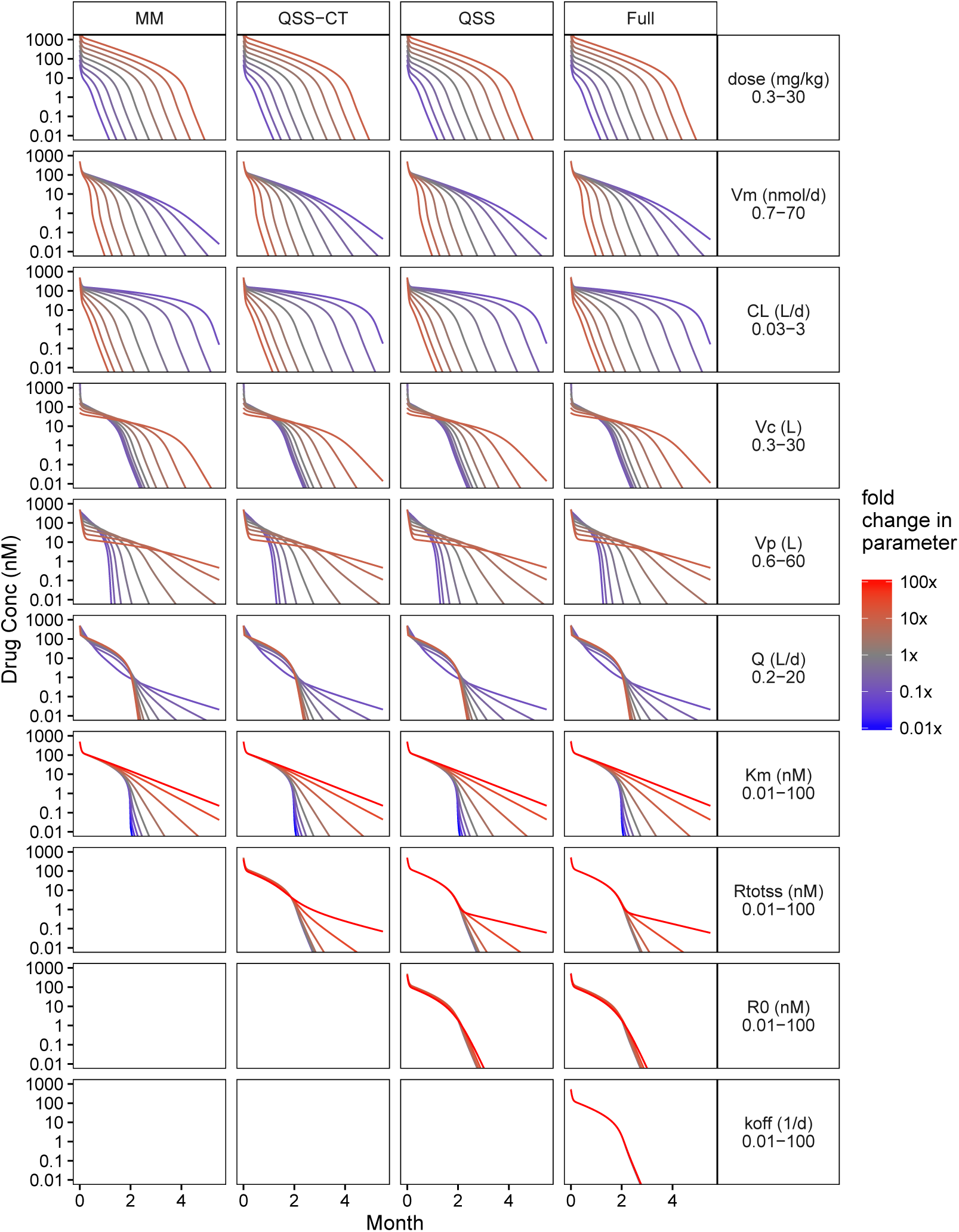
Sensitivity analyses in drug concentration profiles for four different models (MM = Michaelis-Menten approximation, QSS-CT = quasi-steady-state with constant turnover, QSS = quasi-steady state, and Full = Full Model). In this analysis, one parameter is changed throughout the range shown in the y-label on the right, while all the rest are held fixed. Parameters {Dose, *V*_m_, *CL, V*_c_, *V*_p_, *Q*} were varied from 0.1x-10x (by increments 3.16x) from their baseline value while parameters {*K*_m_, *R*_tot,ss_, *R*_0_, *k*_off_} were varied from 0.01x-100x (by increments of 3.16x). Note that varying *R*_tot,ss_ and *R*_0_ while keeping *K*_m_ fixed had almost no effect on the PK profiles. Parameters used in sensitivity analysis were: dose = 3 mg/kg (which is 210 mg for a 70 kg patient or 1400 nmol for a 150 kDa drug), *V*_m_ = 7 nmol/*d, V*_c_ = 3 L, *V*_p_ = 6 L, *CL* = 0.3 L/d, *Q* = 2 L/d, *K*_m_ = 1 nM, *R*_tot,ss_ = 1 nM, *R*_0_ = 1 nM, *k*_off_ = 1/d. The other rate constants, calculated from these lumped constants were *k*_syn_ = 2.3 nM/d, *k*_eR_ = *k*_eCR_ = 24/d, *k*_on_ = 25/(nM · d), *K*_d_ = *k*_off_/*k*_on_ = 0.04 nM.

### Model fitting and likelihood profiling

To assess the identifiability of the TMDD model, the data for mAb#7 was digitized from [11] and the data for mavrilimumab and romosozumab was digitized from [10], noting that the romosozumab figure (Fig 9 in [10]) contained an error in that the doses tested were 1 mg and 5 mg [15] rather than 1 mg and 10 mg.

Monolix 4.3.2 was used to fit the quasi-steady-state constant turnover (QSS-CT) model, which makes the quasi-steady-state approximation (*K*_ss_ = (*k*_off_ + *k*_eCR_)/*k*_on_) and the assumption of constant receptor turnover (*k*_eR_ = *k*_eCR_). Only mean data was fit (rather than population data) and no random effects were included in the model. Thus Monolix used the Nelder-Mead simplex method (using the Matlab fminsearch.m function) for finding the best-fit parameters, rather than Stochastic Approximation Expectation Maximization (SAEM).

To characterize the practical identifiability of the model parameters, a likelihood profiling analysis was done where the receptor density (*R*_0_) was fixed at various values, and the model was refit to the data, as described by Raue et al [5]. The change in −2 × (Log Likelihood) was reported.

## Results

### Sensitivity analysis

The sensitivity analysis is shown in Figure 2. Each column of plots shows a different model approximation and each row shows a set of PK profiles as one parameter is varied while the rest are held fixed. The top 7 rows of plots show dose, the linear PK parameters and the Michaelis-Menten parameters. These plots show large changes in the PK profiles when the parameter is varied over a 100-fold range. Changing these parameters have the same effect on all models explored.

The next row for *R*_tot,ss_ shows that for large steady state receptor densities (30-100 nM), a distinct terminal elimination phase appeared, though note that this phase was not observed in the case studies presented in Figure 3, where the upper bound for the receptor density was around 10 nM (see below). The final two rows are for parameters *R*_0_, and *k*_off_. These parameters have almost no impact on the PK profiles for even a 10,000-fold change in parameters.

**Figure 3:**
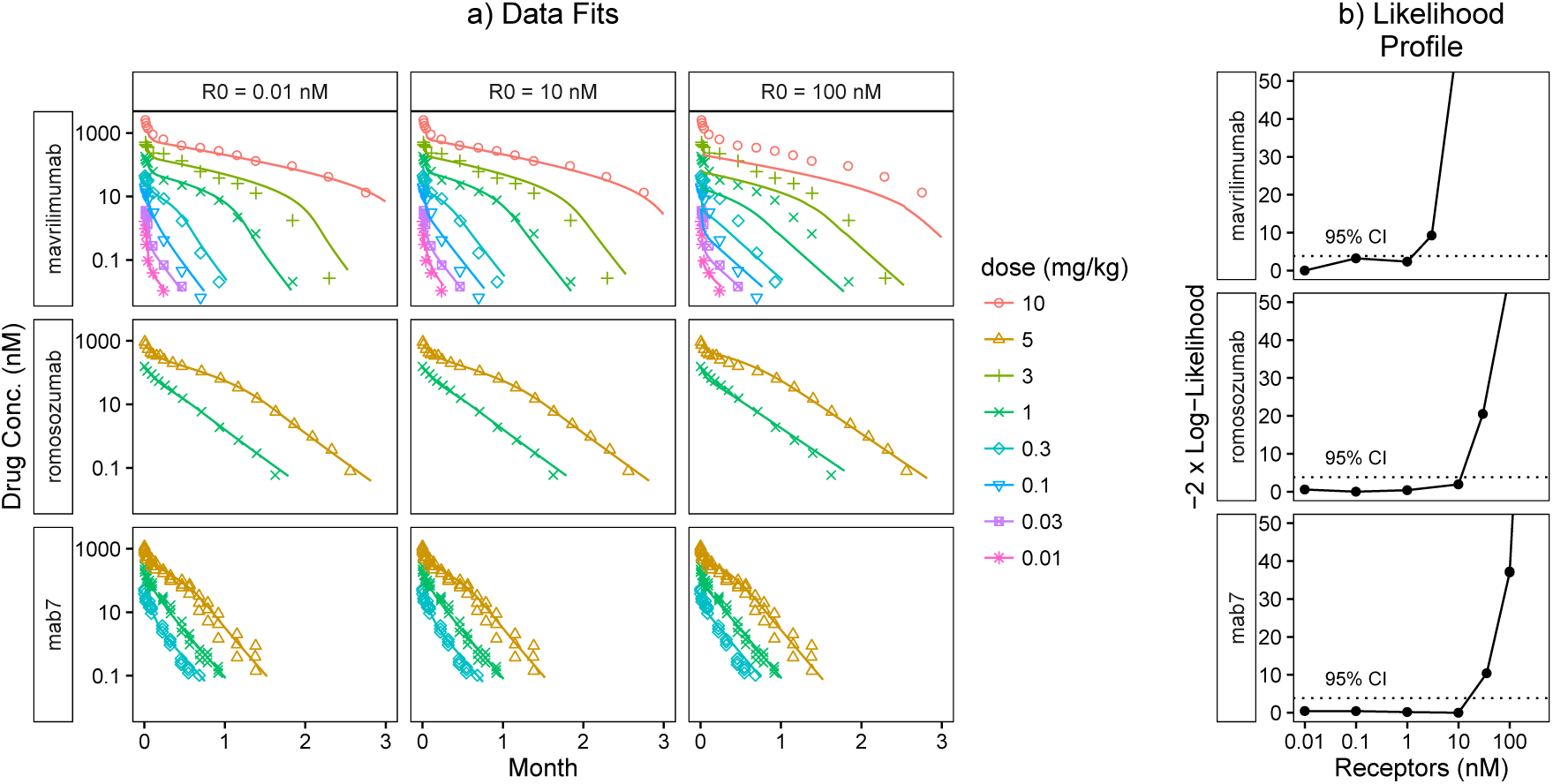
Pharmacokinetics and Likelihood Profiles for mavrilimumab (anti-GM-CSF) [16] and romosozumab (anti-sclerostin) in humans [15] and for un-specified mAb#7 in cynomolgus monkeys [11]. For clarity, a subcutaneous dose of mAb#7 is excluded from the plot, but it was included in all model fitting. Note that for all three drugs, the confidence interval is one sided; there is an upper bound for the receptor number (about 10 nM) but the lower bound is zero. Thus for these datasets, the PK data alone was not sufficient to estimate the receptor density. The 95% Confidence Interval (CI) line was drawn at 3.84, based on a *χ*^2^ distribution with one degree of freedom.

The key insight is that when describing TMDD when only nonlinear PK data is available, there are two key parameters that characterize the profiles: *V*_m_ = *k*_syn_ · *V*_c_ and *K*_m_ = *K*_ss_ · *R*_tot,ss_/*R*_0_. If a terminal phase is detectable, then *R*_tot,ss_ may also be important. For the one-compartment TMDD model (i.e. no peripheral compartment), it has previously been shown that the rate of decline of the terminal elimination is equal to *k*_eCR_ = *k*_syn_/*R*_tot,ss_ [6]. However, the mathematical analysis has not yet been extended to the two-compartment model scenario.

### Model fitting and likelihood profiling

The results of the model fits and likelihood profiling are shown in Figure 3 and the parameters are summarized in Table 2. Note that in the likelihood profiles for all three drugs explored, the lower bound on the receptor density is zero, making the parameter practically identifiable. This can also be observed in the fits to the data where changing *R*_0_ by 1000-fold (from 0.01 nM to 10 nM) has almost no impact on the individual fits.

**Table 2:** Parameter fit of equilibrium constant turnover model to digitized data

|  | mavrilimumab | romosozumab | mab7 | units |
| --- | --- | --- | --- | --- |
| target | GM-CSF | sclerostin | unknown |  |
| species | human | human | cyno |  |
| F | - | - | 0.66 (4.9%) | - |
| ka | - | - | 0.23 (12%) | 1/d |
| Vc | 2.8 (17%) | 2.4 (0%) | 0.16 (6.3%) | L |
| Vp | 5.6 (17%) | 2.6 (3.5%) | 0.1 (6.3%) | L |
| CL | 0.3 (9.6%) | 0.24 (4.4%) | 0.019 (8.8%) | L/d |
| Q | 1.7 (12%) | 0.54 (3.6%) | 0.031 (6.7%) | L/d |
| ksyn | 2.4 (18%) | 6.1 (9.4%) | 16 (9.8%) | nM/d |
| Kss | 1.1 (2.2%) | 12 (6.6%) | 26 (12%) | nM |
| R0 | 0.0064 (110%) | 0.007 (52%) | 0.0036 (23%) | nM |
| keR | 370 | 860 | 4400 | 1/d |

In general, for likelihood profiling analyses, as one parameter is changed, the other parameters may change as well. In this case, for *R*_0_ ≤ 10 nM, the other model parameters did not change significantly. This can also be seen from the QSS-CT sensitivity analysis in Figure 2 which showed that for the constant turnover model, when *R*_0_ = *R*_tot,ss_ ≤ 10 nM, there was almost no impact on the PK profile when changing the receptor density while keeping all other parameters fixed.

## Discussion

The implications of these results are that when a Michaelis-Menten model can adequately describe the available PK data (as is often the case for membrane-bound targets) then PK data alone is not sufficient for the receptor density to be identified. In [10], the authors estimate *R*_0_ = 19 nM for romosozumab which was much higher than the experimental measurements of 0.05 nM. This led the authors to conclude that “low plasma sclerostin concentrations cannot adequately support a large nonlinear clearance if sclerostin-mediated elimination is the reason for non-linear PK.” The authors then concluded that it was more likely that most of the target was in tissue, rather than in the central compartment. However, as shown in Figure 3, the 19 nM estimate is at best an upper bound and low receptor densities of 0.05 nM and below are also consistent with the available data and can explain the PK nonlinearity. Even though the authors used a more complex minimal-PBPK model to describe the data, these same results regarding the unidentifiability of *R*_0_ are expected to apply. In [11], the authors use estimates for *R*_0_ based on model estimates of cynomolgus monkey data to translate across species (assuming that *R*_0_ does not change across species). But as shown here, *R*_0_ is not estimable, and what this analysis highlights is that to translate the TMDD model across species, it is most important to understand how two parameters scale across species: *V*_m_ and *K*_m_ which can be related to the parameters of the TMDD model. One might assume that rate constants scale by weight to the −0.25 power (*k*_eCR_ ~ *WT*^−0.25^), and absent other knowledge, one can assume that receptor concentrations and the quasi-steady-state equilibrium constant (*R*_tot,ss_, *R*_0_, *K*_ss_) is the same across species. Assuming standard allometric scaling (*C*L ~ *WT*^0.75^ and *V* ~ *WT*^1^) gives the scaling below, which was also used by Dong et al. [17].

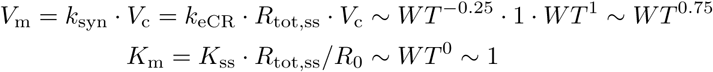

Gibiansky et al., [8, 18] proposed an algorithm where by the full, QSS, MM, and rapid binding model (which uses the assumption that the drug, target, and complex are in a quasi-equilibrium) are all fit to the data and that one then selects “the simplest model that provides predictions sufficiently similar to the predictions of the full model in the dosing/concentration range of interest.” The analysis presented here further supports this recommendation. In particular, in the examples presented here, when receptor concentration data is not available (as is often the case for membrane-bound targets) and when the Michaelis-Menten model is sufficient to describe the data, then the Michaelis-Menten model should be used and estimates for the receptor density should not be made, though it is possible to identify an upper bound for the receptor density.

It is also important to remember that the TMDD model analyzed here is a simplification that leaves out many physiological processes.

1. Target synthesis and distribution in both peripheral tissue [3] and the target tissue (e.g. the joint or tumor) [19, 20].
2. Competition for target binding sites between the drug and the target’s endogenous ligand [21].
3. Feedback mechanisms [22] leading to either an increased synthesis of the target in the presence of drug [23] or a decrease in target synthesis or expression [3, 24].
4. A drug binding multiple targets; the current model assumes implicitly that all the target is measured, but for infliximab binding TNF-*α*, target exists in both membrane-bound and soluble forms, and a model that only accounts for soluble TNF-*α* may considerably underestimate the binding affinity [25].

Because the full TMDD model is a simplification of the true system, even in scenarios when the receptor density is estimable, it will not necessarily be a reflection of the receptor density in the serum. Rather it is a lumped parameter that may contain information about the receptor density in all tissues. This is analogous to how the central and peripheral volumes of two compartment PK models do not necessarily represent physiological volumes [26, chapter 4].

If the modeler desired information about the receptor density, the most straight forward way would be to develop an assay to measure it directly. Alternatively, a highly sensitive PK assay that allows for observation of the terminal elimination phase could help inform the *R*_tot,ss_ parameter and a sensitive assay together with rapid PK sampling could be used to identify *R*_0_, as described previously [6]. However, as the terminal elimination phase occurs at very low drug concentrations, models that accurately describe this portion of the PK profile may not be needed in guiding drug development of the compound.

## Conclusions

The key insight from this analysis is that when fitting a model to PK data for biologics with membrane-bound targets, often a Michaelis-Menten model is sufficient to describe the data. In that scenario, the receptor density is practically unidentifiable and only an upper bound for the receptor density can be identified. An additional insight is that it is the lumped parameter *K*_m_ = *K*_ss_ · *R*_tot,ss_/*R*_0_ that is identifiable and the assumption that is frequently made of constant receptor turnover (*k*_eR_ = *k*_eCR_) is not needed for model fitting and interpretation of the Michaelis-Menten approximation of the full target mediated drug disposition model. Thus even though the full TMDD model is structurally identifiable, it is almost never the case that sufficient data is available for all parameters to be practically identifiable, both because of coarse PK sampling and because of the limit of quantification of the PK assay. The standard confidence intervals reported by Nonlinear Mixed Effect (NLME) modeling software cannot always be used to assess this issue as they are often based on a linear, asymptotic approximations in the neighborhood of the optimal set of parameters for describing the data.

Thus we recommend that pharmacometricians regularly use bootstrapping or likelihood profiling [5] as an additional check. New methods for assessing model parameter uncertainty may also be useful in assessing this issue [27, 28]. This analysis further supports the advice from Gibiansky et al. [8], to fit the Michaelis-Menten (MM), QSS (with constant turnover) and the full model to the data and then to only draw inferences and make predictions using the simplest model that describes the data well.

## Acknowledgements

The author would like to thank Kostas Biliouris, Carter Cao, Alison Margolskee, Phil Lowe, Prasad Ramakrishna, Pratap Singh, and Jean-Louis Steimer for helpful discussions and review of this document.

